# STRIDE: A Sequencing Depth-Insensitive Metric for Robust Comparison between Sparse Chromosome Conformation Capture Data

**DOI:** 10.1101/2025.10.08.681271

**Authors:** Bingxiang Xu, Xiaomeng Gao, Weichu Tao, Zhihua Zhang, Feifei Li

## Abstract

The study of three-dimensional (3D) genome organization has been revolutionized by high-throughput chromatin conformation capture technologies (Hi-C) or its derivatives. However, the dependency on sequencing depth severely restricts the reliability and accuracy of existing Hi-C tools, especially in single-cells. To address this issue, we introduce a novel computational framework based on the mean first passage time (MFPT) in Markov chain theory, which transforms chromatin contact matrices into a robust, distance-based representation. We demonstrate that MFPT representation is inherently insensitive to sequencing depth. Leveraging this transformation, we develop STRIDE (Spatial Topological Representation of Interaction Distance Evaluation), a parameter-free metric for library similarity. STRIDE is resilience to sparse and noisy data, insensitive to technical variabilities, and facilitates unsupervised embedding from single-cell Hi-C, enabling accurate delineation of cellular states and developmental trajectories. In conclusion, as a reliable computational framework for sparse and noisy data, STRIDE may serve as a base for wide range single-cell 3D genome analysis that used to be inconceivable.

## Introduction

High-throughput chromatin conformation capture techniques, including Hi-C, ChIA-PET, and HiChIP/PLAC-Seq, have significantly advanced our understanding of 3D genome structure by enabling high-resolution, genome-wide mapping of chromosomal interactions ^1–4^. These methods facilitate analysis across multiple scales—from large A/B compartments to smaller TADs and chromatin loops—and reveal dynamic changes in 3D genome organization during development and environmental responses ^5–11^.

Hi-C data’s two-dimensional structure necessitates quadratically scaled sequencing to achieve high-resolution maps ^12,13^. For example, generating 1Kb resolution contact maps with adequate quality in human GM12878 cells requires ∼5 billion sequenced reads ^14^. This sequencing depth is often impractical, especially at the single-cell level, where only 5k–100k read pairs are typically obtained per cell ^15–17^. As a result, most Hi-C studies suffer from insufficient or unsaturated sequencing depth, compromising data reliability and sensitivity and hindering direct comparisons between libraries ^16^.

Library consistency or distance comparison is a fundamental requirement in Hi-C data analysis. Various strategies have been seen in literature, including smoothing via local averaging (e.g., HiCRep ^18^ and QuaSAR-seq ^19^), spectral decomposition (HiC-spector ^20^), and missing value imputation with noise reduction using random walk models (genomeDISCO ^21^). Those methods primarily aim to mitigate experimental noise and avoid overestimating library consistency due to the high correlation between contact frequencies and genomic distances. However, those random down-sampling or standardize library depth-based methods subjected to severe wastes of valuable data and risks in additional stochastic fluctuation.

Emerging evidences indicated that contact frequencies are inherently correlated ^1,22^, especially between adjacent or spatially proximal regions ^23,24^.

These correlations offer valuable insights into chromatin conformation and provide an opportunity to reduce reliance on sequencing depth for data representation. Leveraging these correlations could enable a more robust and cost-effective framework for chromatin conformation mapping.

Based on these insights, we introduce a novel computational framework that uses the mean first passage time (MFPT) to derive a robust, distance-based representation of chromatin contact data. MFPT calculations leverage correlations between proximal bins and are inherently insensitive to sequencing depth variations. Building on this, we developed STRIDE (Spatial Topological Representation of Interaction Distance Evaluation), a parameter-free metric for library (dis)similarity that eliminates hyperparameter tuning. We demonstrate that STRIDE is robust to sequencing depth fluctuations and technical noise while improving the detection of biologically meaningful differences. Additionally, STRIDE facilitates unsupervised embedding of single-cell Hi-C data, enabling its application at the single-cell level.

## Results

### MFPT matrices stably preserve structural details in Hi-C data in reduced sequencing depths

The STRIDE algorithm leverages the sequencing depth-insensitive properties of mean first passage time (MFPT) representation derived from contact matrices. Unlike contact frequency-based approaches, MFPT quantifies the distance or separation between bin pairs by calculating the average number of transitions required to move between bins via random walks, where contact frequencies define transition probabilities. Specifically, a balanced Hi-C contact matrix is interpreted as a transition matrix for a finite-state Markov chain (random walk process). For irreducible and positive recurrent chains, the MFPT between bins reflects their separation, with transitions dominated by high-probability paths (strong contacts), which remain stable under reduced sequencing depth. This confers two key advantages: (1) reduced sensitivity to sequencing depth, as low-probability transitions (weak contacts) minimally impact MFPT values, and (2) preservation of spatial information for bin pairs with zero observed contacts, as MFPT incorporates indirect connectivity via intermediate bins. These features enable robust inference of chromatin organization even with sparse data. Details are provided in the Methods section.

To evaluate the sequencing depth insensitivity of MFPT representation, we tested it using a series of down-sampled datasets from the deeply sequenced GM12878 cells at sampling rates of 1/2, 1/4, 1/8, 1/16, and 1/32 (Fig. 1A) ^14^. The lowest-depth library (1/32) contained ∼80 million valid cis contact pairs (genome distance ≥20 kb), with 72.5% of bin pairs exhibiting zero interactions and 66% of annotated loops (10 kb resolution) lost, resembling about pooled ∼1000–2000 single-cell Hi-C data ^25^. Globally, contact map noise increased rapidly with reduced sequencing depth, obscuring topological features (Fig. 1A), while MFPT maps largely retained their original structure even at 1/32 depth (Fig. 1B). This robustness persisted in local regions near diagonals of the contact maps, where read densities were higher (Fig. 1A, B). Quantitatively, we calculated a signal-to-noise ratio (SNR) for each euchromosome to assess noise introduced by down-sampling (Supplementary Text). MFPT maps consistently achieved higher SNR across all chromosomes and sequencing depths (Fig. 1C).

**Fig. 1.**
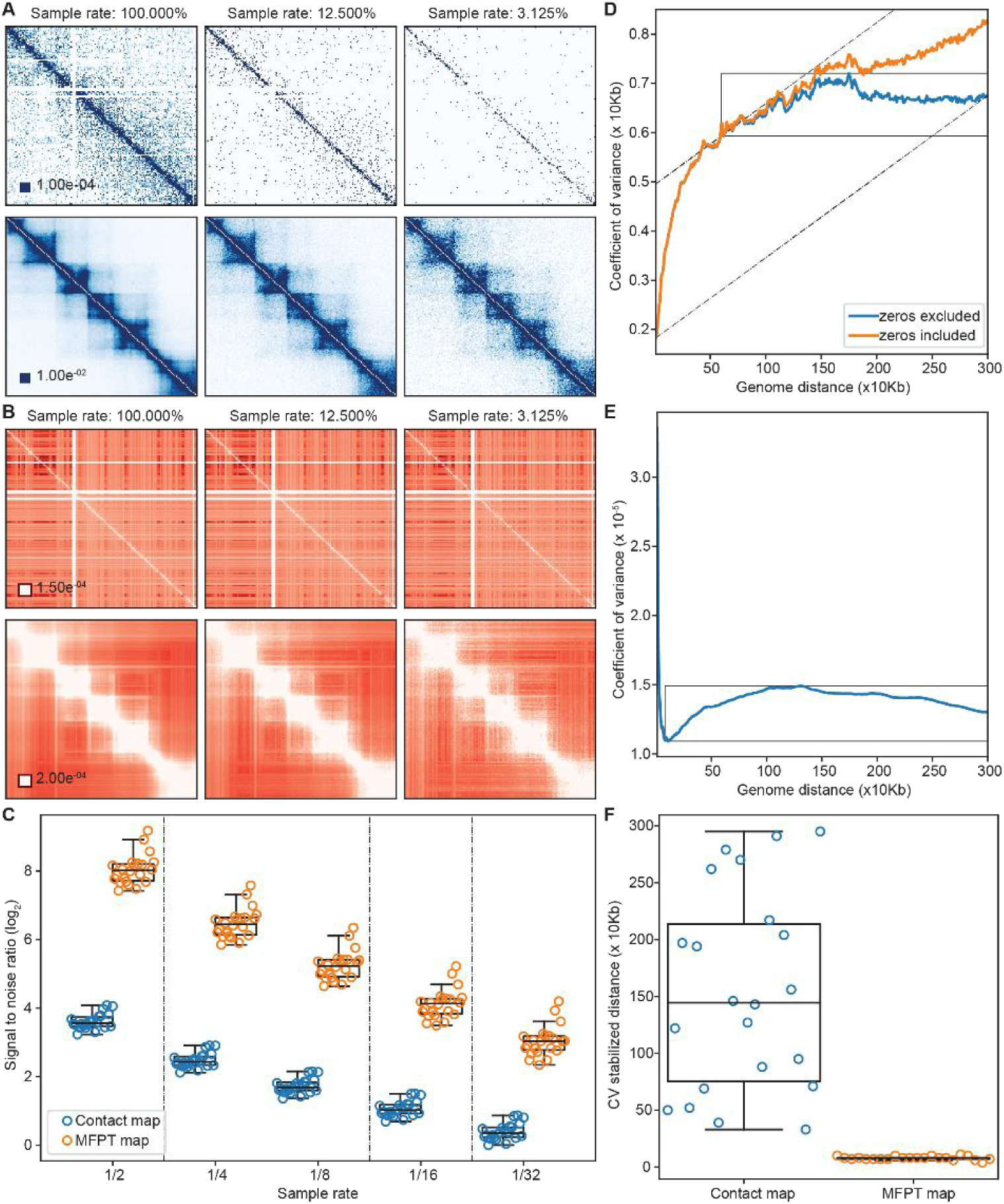
MFPT matrices retain structural features of Hi-C data under lower sequencing depths. **(A)** and **(B).** Contact maps (A) and heatmap of MFPT matrices (B) for chromosome 6 from different levels of down-sampled libraries of the GM12878 dataset. A uniform resolution of 10 kb was used. The entire chromosome is displayed in the top row and only one out of every ten bins is displayed to enhance contrast. The bottom row shows the local region of chr6: 42.00 - 44.00 Mb. **(C)** The signal to noise ratio values of contact maps and MFPT maps at local regions. Chromosome 1 to 22 were shown from left to right in each sequencing depth. **(D)** and **(E).** The relationship between the variability (coefficient of variation, CV) of contact frequencies (D) or MFPT values (E) and genomic distances is shown. Chromosome 6 is shown as an example. Rectangular boxes highlight the regions where CV values stabilized. In panel (D) it was calculated using contact matrix excluding frequencies of zero (blue line), while yellow line (including frequencies of zero) did not reach stabilization. Two dashed lines in panel (D) indicate the calculation of genomic distance at which CV values reached stabilization. **(F).** The distribution of chromosome wise genomic distances at which CV values stabilized in contact matrices and MFPT maps for all chromosomes.

Next, we showed the weak dependency between MFPT variability and genomic distance (Fig. 1D, E; Extended Data Fig. 1). This insensitivity arises from the MFPT calculation, which accounts for all bin pairs irrespective of genomic distance. By plotting the coefficient of variation (CV) for contact frequencies or MFPTs against their genomic distances across chromosomes in GM12878 data (Fig. 1D, E), we identified the genome distance at which CV stabilized (Supplementary Text). Contact frequency variability increased with genomic distance and remained unstable even after excluding zero-frequency pairs (Extended Data Fig. 1), particularly at long distances (330 kb–2.95 Mb; 690 kb for chromosome 6, Fig. 1D, F). Conversely, MFPT variability decreased with genomic distance and stabilized rapidly at shorter distances (40 kb–140 kb; 70 kb for chromosome 6, Fig. 1E, F).

In summary, MFPT representation is robust to sequencing depth and maintains conformational patterns at low coverage. Critically, MFPT variability exhibits negligible correlation with genomic distance, enabling downstream analyses without normalization for distance-related dimensional distortions.

### MFPT representation preserves topological features in low-depth Hi-C libraries

The MFPT representation accurately retained TAD and loop structures of chromatin organization in down-sampled GM12878 datasets. Using insulation score (IS) to predict TAD boundaries ^26^, MFPT-derived IS profiles consistently localized peaks to annotated TAD boundaries across all sampling depths (Fig. 2A), with periodicity matching established TAD size distributions. Chromosome-wide IS profiles remained highly stable during downsampling, exhibiting pairwise correlations >0.95 between consecutive depths—comparable to contact map-derived profiles. However, quantitative analysis of sequencing depth sensitivity (Supplementary Text) near TAD boundaries revealed critical differences: conventional contact maps showed marked inter-domain contact frequency deterioration at low depths (Extended Data Fig. 2A, B; Supplementary Text), while MFPT maintained more robust inter-domain distances irrespective of sequencing depth coverage (Fig. 2B; Extended Data Fig. 2B).

**Fig. 2.**
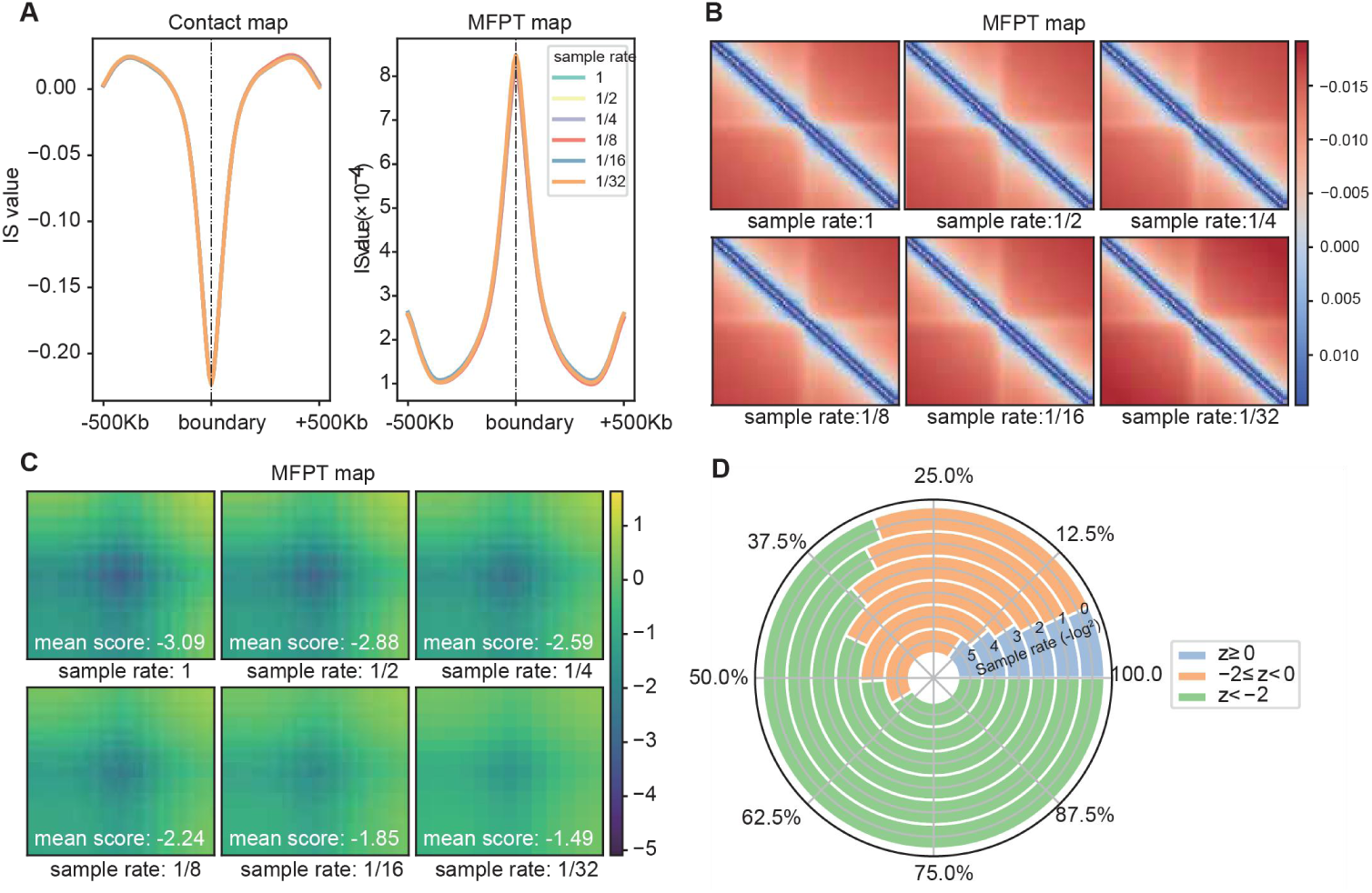
MFPT matrices robustly preserve TAD and loop features in low-coverage Hi-C datasets. **(A)** Stacked insulation score values around annotated TAD boundaries, computed from both contact and MFPT maps at varying sequencing depths. **(B)** Heatmap showing the averaged MFPT profile around annotated TAD boundaries at varying sequencing depths. The top-left and bottom-right corners represent adjacent TADs. **(C)** The APA profiles calculated from the MFPT maps in all sampling rates, with average z scores marked (see methods). **(D)** Proportions of the annotated loops with significantly enriched contact signals (*z* <− 2), enriched but non-significant contact signals (−2 ≤ *z* < 0), no enrichment in contact signals (*z* ≥ 0) based on the scores calculated from MFPT maps.

The MFPT framework preserved enriched contact signals around chromatin loops at low sequencing depths. Both globally (averaging all loops) and locally (single-loop analysis) in GM12878 cells ^27^, the decline in loop enrichment scores (Methods) was more gradual in MFPT maps than in original contact maps (Fig. 2C; Extended Data Fig. 2C–E). Initially, conventional contact maps reported more annotated loops due to their high-coverage origin, whereas MFPT maps underrepresented these loops (Fig. 2D; Extended Data Fig. 2F). However, this discrepancy diminished rapidly with decreasing sequencing depth and reversed at sampling rates below 1/8, demonstrating MFPT’s enhanced robustness in retaining loop-associated signals under sparse coverage, since the loop positions were determined by the contact maps themselves.

In conclusion, MFPT representation effectively preserves topological features of contact maps—such as TAD structures and loop-associated signal enrichment—and provides a robust foundation for downstream Hi-C feature detection under low-coverage conditions.

### STRIDE is robust in library consistency measurement across variable sequencing depths

Leveraging MFPT’s insensitivity to sequencing depth, we developed STRIDE, a novel, sequencing-depth-insensitive metric for library distance (Methods). STRIDE directly computes dissimilarity between libraries by comparing their MFPT representations, obviating preprocessing steps like downsampling. STRIDE was benchmarked against established metrics—HiCRep ^18^, HiC-spector ^20^, and genomeDISCO ^21^ (Methods)—with consistency scores subtracted from 1, so that more similar libraries yield values closer to 0. Critically, it requires no tunable hyperparameters (Methods), ensuring robust and parameter-free performance.

First, STRIDE is insensitive to sequencing depth. Using down-sampled synthetic datasets from GM12878 and K562 cell lines ^14^, at sampling rates of 1/ 2, 1/4, 1/8, 1/16, and 1/32, libraries were categorized into three replicate types—pseudo-replicates (PR), biological replicates (BR), and non-replicates (NR) ^18^. A robust library similarity metric should satisfy PR > BR > NR across all sequencing depths and exhibit decreasing consistency with increasing down sampling rates (due to elevated stochastic noise). STRIDE and genomeDISCO adhered to these principles consistently across all chromosomes, followed by HiCRep, which deviated in two chromosomes (Fig. 3A; Extended Data Fig. 3). HiC-spector, however, failed in 16 chromosomes (Fig. 3A; Extended Data Fig. 3), revealing its inability to mitigate noise from random fluctuations or technical variability. Consequently, HiC-spector was excluded from further analysis in this section.

**Fig. 3.**
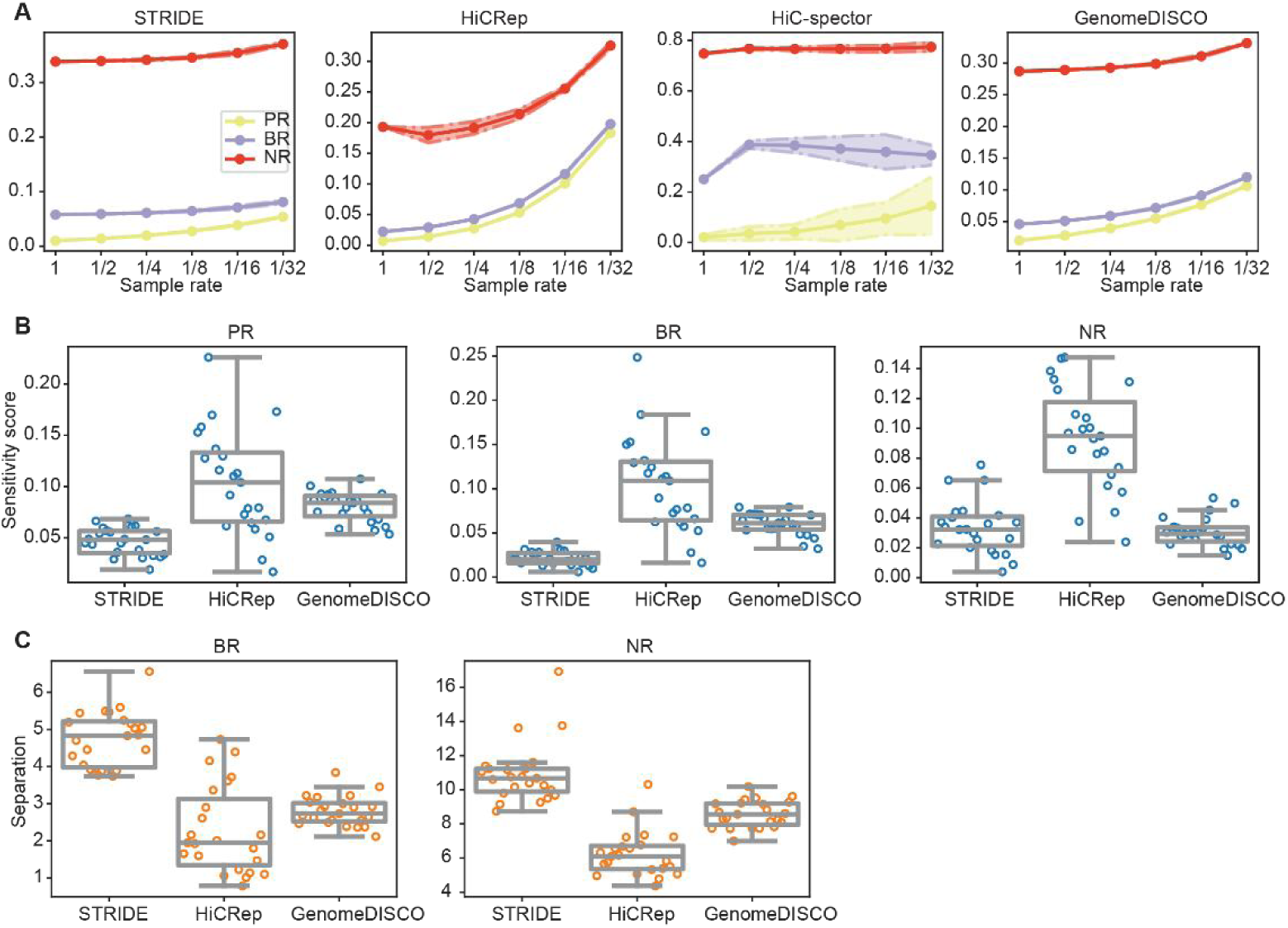
STRIDE demonstrates consistent similarity measurements across varying sequencing depths. **(A)** Dissimilarity scores for pseudo-replicates (PR), biological replicates (BR), and non-replicates (NR) across different sampling rates were calculated using four methods. Chromosome 1 is shown as an example, with shaded areas representing confidence intervals from 10 independent down-sampling trials. **(B)** The distribution of sequencing depth sensitivities (see Methods) is compared across all chromosomes for STRIDE, HiCRep, and genomeDISCO, within PR, BR and NR, respectively. **(C)** The discrimination index (see Methods) between BR and PR is shown on the left, and that between NR and BR on the right, for STRIDE, HiCRep, and genomeDISCO. In panels (B) and (C), chromosomes 1 to 22 and X are represented as dots from left to right.

STRIDE demonstrated the least sensitivity to sequencing depth, evident from its most gradual score changes across down sampling rates (Fig. 3A; Extended Data Fig. 3). Quantitative analyses confirmed STRIDE’s superior stability in comparing PRs and BRs (all with *p* < 10*⁻*³, paired *t*-tests, chromosome-wisely; Fig. 3B; Extended Data Fig. 4A). For NRs, STRIDE outperformed HiCRep (mean difference = 0.060, *p* = 2.8 × 10*⁻⁹*) but aligned closely with genomeDISCO (mean difference = 0.00266, *p* = 0.44; Fig. 3B). Notably, HiCRep remained the most sequencing depth sensitive metric even after down sampling normalization (Methods).

Second, STRIDE is capable to discriminate PRs, BRs, and NRs. Discrimination performance was quantified by the noise level (introduced by random down-sampling) required to degrade PR/BR consistency to BR/NR levels (Supplementary Text; Extended Data Fig. 4B). STRIDE consistently outperformed HiCRep and genomeDISCO in distinguishing PRs vs. BRs and BRs vs. NRs (all with *p* < 10*⁻⁵*, paired *t*-tests; Fig. 3C), underscoring its enhanced robustness in resolving library relationships under noise.

In conclusion, STRIDE exhibited minimal sensitivity to sequencing depth compared to other commonly used Hi-C (in)consistency metrics and achieved optimal discrimination of replicate types (PRs, BRs, NRs), solidifying its utility for robust library comparison under variable sequencing depth coverages.

### STRIDE is insensitive to technical variability in experimental protocols

To assess the sensitivity to technical variability, we analyzed a dataset of 8 Hi-C libraries generated using diverse protocols—including *in situ* versus dilution Hi-C, HindIII versus MboI/DpnII digestion, FA versus FA+DSG crosslinking—in hESC and IMR90 cells (Fig. 4A; Extended Data Table 1 ^18,28–30^). Clustering hierarchies derived from an ideal metric should adhere to the following criteria: 1) cells of the same type should be prioritized for clustering; 2) within the same cell type, libraries use the same experimental protocol should be clustered together; and 3) the lower-quality HindIII-digested hESC library (rep1) ^18^ should maintain a certain distance from other libraries (Extended Data Fig. 5A).

**Fig. 4.**
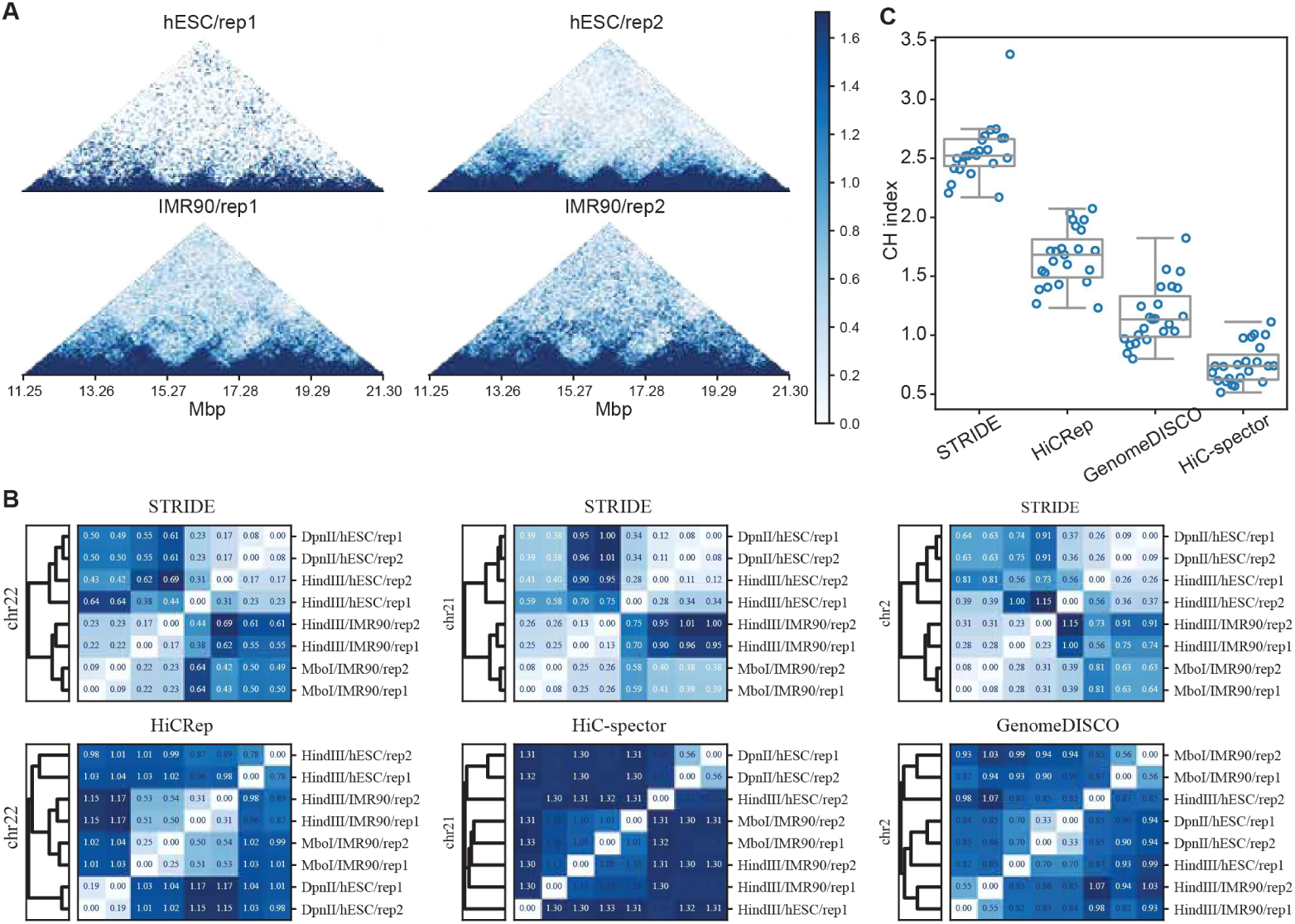
STRIDE exhibits higher tolerance to variations in the experimental protocols. **(A)** Contact maps of Hi-C libraries digested by HindIII from hESC and IMR90 (chrX:11250000-21250000). **(B)** Library distances calculated by each consistency metric and hierarchical clustering results based on these distances, with only some chromosomes shown as examples. **(C)** Chromosome-wise CH index values calculated from each metric for evaluating the separation between the two cell types (chromosomes 1 to 22 and X are arranged from left to right as dots).

STRIDE was the sole method that fully satisfied criteria 1) and 2) across all chromosomes. The only deviation was the low-quality HindIII-digested hESC library misclustered with those digested with DpnII on chromosomes 1, 2, 3, 17, and 19 (Extended Data Fig. 5B). HiCRep incorrectly grouped one HindIII-digested hESC with IMR90 libraries on chromosomes 19, 20, and 22 (Fig. 4B; Extended Data Fig. 4C), while HiC-spector (Fig. 4B; Extended Data Fig. 4D) and genomeDISCO (Fig. 4B; Extended Data Fig. 4E) erroneously merged distinct cell types across many chromosomes. All three alternative metrics also failed to properly distinguish the low-quality HindIII-digested hESC library on multiple chromosomes.

Moreover, STRIDE consistently outperformed the other metrics in quantitatively discriminating cell types across all chromosomes (Fig. 4B; Extended Data Fig. 5). Cell type discrimination was quantified using the Calinski-Harabasz index (CH index), which compares the average inter cluster distance between libraries from different cell types to that within the same cell type. Across all chromosomes, STRIDE achieved a minimum CH index of 2.17 (on chromosome 19), surpassing the maximum CH indices of HiCRep (2.07), HiC-spector (1.11), and genomeDISCO (1.82) (Fig. 4C). This demonstrated STRIDE’s robustness in maintaining cell type-specific clustering under technical variability.

In conclusion, STRIDE exhibits superior tolerance to experimental protocol variations compared to other Hi-C consistency metrics.

### STRIDE accurately infers lineage relationships across cell types in sparse Hi-C data

To benchmark the algorithms’ capacity for cell lineage inference, we applied them to a mouse hematopoiesis dataset comprising ten cell types ^9,31^. The reference cell type phylogeny was adapted from a version derived from an RNA-seq dataset ^32^ (Extended Data Fig. 6A–B). STRIDE accurately discriminated precursor cells from terminally differentiated granulocytes (GR) and megakaryocytes (MK), as well as erythroid from myeloid lineages (Fig. 5A; Extended Data Fig. 7). Using the generalized Robinson-Foulds distance to compare inferred phylogenies with the reference, STRIDE outperformed the other three metrices uniformly on 16 out of 19 euchromosomes (mean RF distance reductions: −0.17 vs. HiC-spector, −0.24 vs. genomeDISCO, −0.31 vs. HiCRep; all with *p* < 2.5 × 10*⁻⁵* via paired *t*-tests; Fig. 5B). Additionally, only STRIDE and genomeDISCO consistently surpassed random phylogenies across all chromosomes (20,000 trials, *p* < 0.05).

**Fig. 5.**
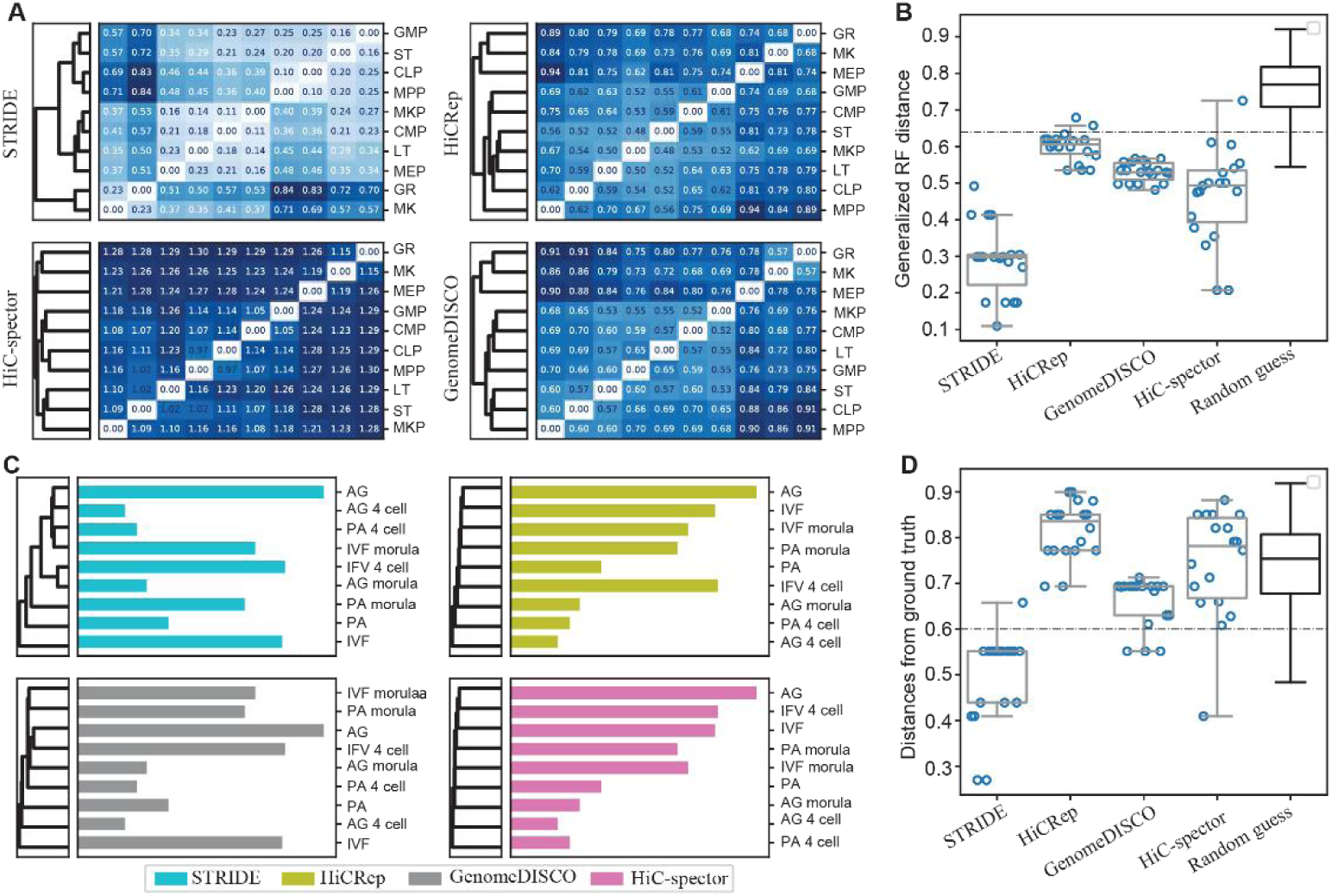
STRIDE infers lineage relationships across diverse cell types. **(A)** Library distances between cell types in mouse hematopoietic system calculated by each method and hierarchical clustering of cells based on the calculated distances, with chromosome 5 shown as example. **(B)** Chromosome wise generalized RF distances between phylogenic relationships derived from different methods and the ground truth (Figure S5B) in the mouse hematopoietic system. Chromosomes 1–19 were displayed as dots from left to right. “Random guess” represented the theoretical distribution of generalized RF distance calculated from 20,000 random phylogenies. **(C)** Hierarchical clustering of cell types during porcine early embryonic development based on the libraries’ distances calculated by each method, with chromosome 3 shown as an example (left panel). Widths of the bars indicated sequencing depth for corresponding libraries (right panel). **(D)** Same as (B) but for cell types in porcine early embryo development. Abbreviations: long-term/short-term hematopoietic stem cells (LT/ST-HSCs), multipotent progenitor (MPP), common myeloid progenitor (CMP), common lymphoid progenitor (CLP), granulocyte-macrophage progenitor (GMP), megakaryocyte-erythrocyte progenitor (MEP), megakaryocyte progenitor (MKP), granulocytes (GR) and megakaryocytes (MK), in vitro fertilization (IVF), parthenogenesis (PA) and androgenesis (AG).

Next, we assessed the four metrics in cell lineage inference using a low sequencing depth Hi-C dataset from early porcine embryo development (182–587 million read pairs per library ^33^, Fig. 5C–D). STRIDE again demonstrated superior performance, accurately capturing the rapid chromatin remodeling in maternal chromosomes compared to paternal ones (Extended Data Fig. 6C; Fig. 5C; Extended Data Fig. 8). Notably, phylogenies provided by HiCRep and HiC-spector were dominated by sequencing depth in many chromosomes, whereas STRIDE remained unbiased (Fig. 5C; Extended Data Fig. 8), reinforcing its robustness to technical limitations.

In conclusion, STRIDE uniquely balances resilience to experimental noise with the preservation of biologically meaningful library differences, enabling more accurate cell lineage inference.

### STRIDE facilitates unsupervised embedding of single-cell Hi-C data

STRIDE can resolve intercellular lineage relationships with single-cell Hi-C data. We demonstrate this with two datasets: the mESC cell cycle ^10^ dataset (1,572 cells) and the mouse cortical / hippocampal neuron ^11^ dataset (1,954 cells). To isolate STRIDE’s contribution, we adopted a minimal downstream pipeline—directly applying tSNE to STRIDE distance matrices without additional imputation or enhancement. The visualization based on tSNE dimensional reduction clearly revealed biologically meaningful clustering. In the cell cycle dataset, large clusters of cells aligned with the temporal order of phases (G1, early-S, late-S/G2), with late-S/G2 cells partitioned into two subgroups, likely reflecting late-S and G2 stages (Fig. 6A; Extended Data Fig. 9). In the mouse neuron dataset, the cells were generally positioned along the order of neurons, oligodendrocytes, astrocytes, and microglia (Fig. 6B and Extended Data Fig. 10). A clear boundary existed between cells originating from the neuroectoderm (neurons, oligodendrocytes, astrocytes) and those originating from the mesoderm (microglia) ^34^. Additionally, there was partial overlap between oligodendrocytes and astrocytes, consistent with the fact that both of them partially originate from the gliogenic progenitor cells ^35^.

**Fig. 6.**
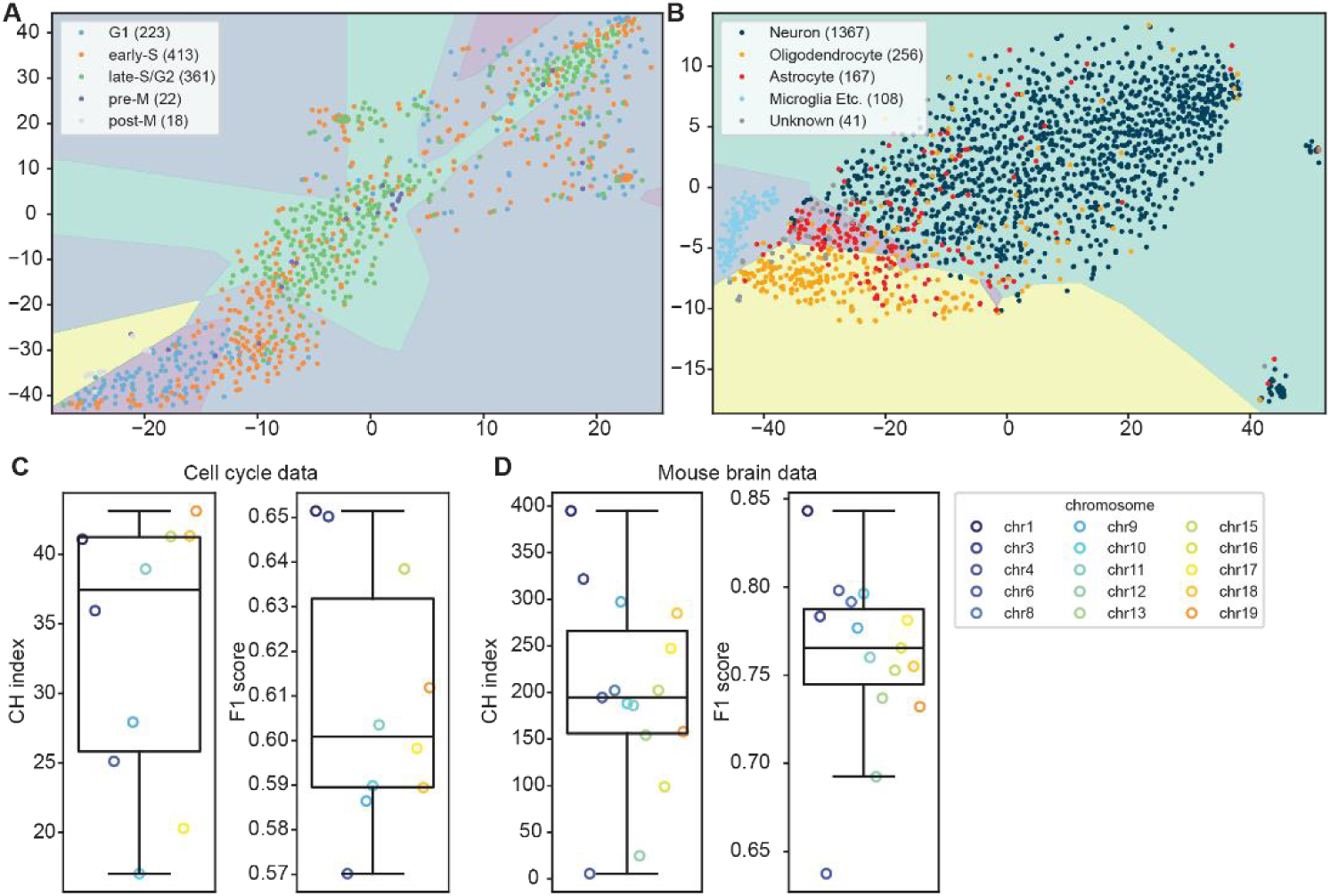
STRIDE facilitates unsupervised dimensionality reduction and embedding of single-cell Hi-C data. (A and. **B)** The t-SNE low-dimensional embedding results for the cell cycle (A) and mouse neuron (B) datasets. Chromosome 1 (A) and chromosome 10 (B) are shown as representative examples. The number of cells for each corresponding cell type is indicated in parentheses. Shaded areas represent cell type separation results based on the MLP model. **(C and D)** Quantitative assessment of cell type classification performance using CH index and F1 score in the cell cycle (C) and mouse neuron (D) datasets.

Quantitatively, the CH index, computed from tSNE analyses utilizing established cell type annotations were uniformly larger than 1 across all chromosomes (Fig. 6C and 6D). This observation affirmed the discernible separation among distinct cell types as inferred from cell-to-cell distances calculated via STRIDE. Furthermore, the tSNE-derived spatial organization of different cell types proved amenable to partitioning through rudimentary machine learning algorithms. In the context of the cell cycle dataset, a three-layer MLP architecture with a configuration of 16 × 16 × 16 neurons achieved division accuracies approximating or surpassing 60% on all scrutinized chromosomes, despite the paucity of cells in the M phase and the confounding amalgamation of late-S/G2 cells (minimum weighted average F1 score = 0.57, Fig. 6C). Conversely, in the mouse neuron dataset, a more parsimonious three-layer MLP model, characterized by an 8 × 8 × 8 neuronal arrangement, sufficed to elevate division accuracy to 75% or greater on the preponderance of chromosomes, with the exception of chromosome 4, wherein the model inadvertently consolidated astrocytes and oligodendrocytes into a single classificatory category (Fig. 6D; Extended Data Fig. 10).

It has been documented that integration of HiCRep with Multidimensional Scaling (MDS) also yielded commendable embedding outcomes ^36^. However, these pairwise-based methods were computationally expensive when used to large numbers of cells. In contrast, STRIDE’s design allows efficient batch processing (Methods). On a standard workstation, STRIDE can compute a distance matrix for around 2000 cells in just seconds—dramatically faster than HiCRep, which takes hundreds of hours.

In conclusion, STRIDE can efficiently and rapidly calculate the distances among the large number of cells and multiple cell types in single cell Hi-C data.

## Discussions

In this study, we adapted the mean first passage time (MFPT) concept from Markov chain theory to model high-throughput chromatin conformation capture data, established a novel MFPT representation framework. This transformation is inherently insensitive to sequencing depth variations, and faithfully preserves topological information embedded in contact matrices. Building upon these advantageous properties, we developed a robust metric termed STRIDE, which is tolerant to sequencing depth, technical noise, and experimental protocols, for assessing library consistency and inter-library distances. Notably, STRIDE maintains these advantages even when applied to the challenging single-cell Hi-C data.

The MFPT representation diverges from conventional Hi-C data analysis methods in three ways ^23^. Firstly, the MFPT systematically captures and utilizes the inherent contact frequency dependencies of neighboring genomic loci, thereby preserving the structural information manifested as correlated contact patterns ^37^. Secondly, MFPT effectively fills in the missing values in contact maps in conditions of insufficient sequencing depth, thereby preserving the topological features of the contact maps. Thirdly, the variation of MFPTs shows significantly weaker correlation with genomic distance compared to contact frequencies, making MFPT-based modeling much simpler and eliminating the need for further normalization. Thus, MFPT pave the way for the development of future bioinformatic tools that maintain robust performance even under low-coverage sequenced conditions.

MFPT’s advantage stems from three fundamental theoretical merits: (1) Its mathematical existence, stability, and computability are rigorously guaranteed by the well-established theory of mean first passage time in irreducible and positive recurrent Markov chains, offering a robust theoretical foundation; (2) Unlike model-dependent approaches, it does not require assumptions about prior probability distributions or complex neural network architectures, thereby eliminating the need for hyperparameter tuning entirely; (3) As a self-contained transform, its computation relies solely on intrinsic information from the input contact matrix, with no external training data dependency. These inherent properties of the MFPT representation contribute to STRIDE’s robustness in handling low-coverage datasets, consistent performance across diverse experimental protocols, and reliable preservation of biologically relevant variations in chromatin architecture.

The limitations of MFPT mainly settled in the following two. First, when sequencing depth is adequate, the MFPT representation’s characteristic of considering all contact frequencies in computing distances between any loci results in over-smoothing of the data, thereby diminishing the contrast in the maps. This may reduce the signal strength of structural elements at higher sequencing depths. Second, the computation of the MFPT representation disrupts the natural sparsity of the contact map, with storage requirements being proportional to the square of the number of bins and non-compressible. This may present challenges for computing MFPT at high resolutions. However, sequencing depths for most of Hi-C studies are far from saturated, especially in the context of single-cell omics, where data sparsity remains a major challenge. Therefore, we expect that MFPT will be applicable to the vast majority of practical scenarios.

Single-cell and spatial Hi-C technologies are becoming key tools for studying 3D genome organization and its functional roles in development and disease. The sequencing depth achievable from one or a few cells remains extremely limited ^16,38^. In this context, MFPT emerges as a theoretically robust analytical method capable of efficiently capturing the topological features of chromatin conformation and identifying relatively homogeneous subgroups within heterogeneous cell populations. The MFPT representation is expected to find broad applications, such as uncovering tumor heterogeneity and tracking cell fate decision trajectories.

## Methods

### Calculation of MFPT representation

The calculation of MFPT representation relies on the theory of the mean first passage time in Markon chain. Consider a (diagonal removed, intrachromosomal) contact matrix *c* = {*c_𝑖j_*}_n×n_, where c_𝑖j_ represents the number of contact products between bins *i* and *j* and, all *c_ii_* set to 0. Clearly, *C* is a symmetric matrix with all elements nonnegative. Through the matrix balancing normalization such as ICE and KR, matrix *C* can be easily normalized into a matrix *p* = {*p_𝑖j_*}_n×n_, satisfying 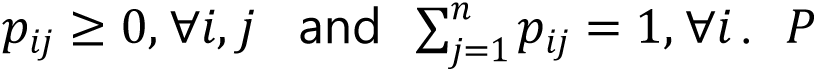 can simultaneously be viewed as the adjacency matrix of an undirected graph and the probability transition matrix of a Markov chain (random walk) on this graph. It is noted that, as long as 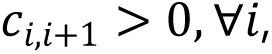 that is, the sequencing depth is sufficient to capture at least one contact product from any bin to its neighbors, the undirected graph corresponding to *P* is connected, and the corresponding random walk process is irreducible and positive recurrent. Let the path of this random walk process be 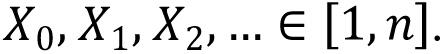 The first passage time from bin *i* to bin *j* is defined as

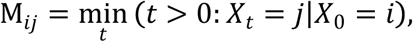

which represents the number of steps required for the random walk process to reach state *j* at first time when starting from state *i*. The mean first passage time (MFPT) from *i* to *j* is then defined simply as

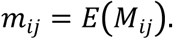

By the properties of conditional expectation and the memoryless nature of Markov chains, it is straightforward to establish the following fact:

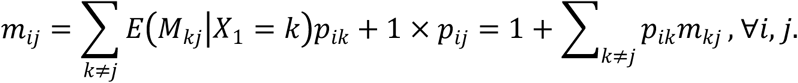

or expressed as the following matrix equation:

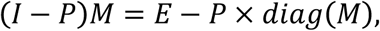

where 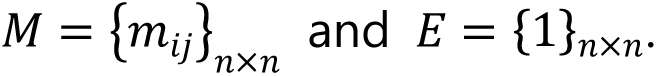

The solution to this equation has been well-studied ^39^, and one common method involves introducing the following auxiliary matrix

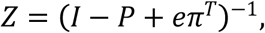

where *I* is the identity matrix, *π* represents the stationary distribution of the random walk process and *e* = {1,1, ⋯, 1}*^T^* is a vector with all elements 1. Then it has been shown that

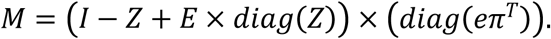

Since *P* is a symmetric probability transition matrix, it is evident that *e* satisfies the equation for the stationary distribution, *e* × *P* = *e*. Therefore, it follows that *π* = *e*/*n*. Thus, the expression for Z can be simplified to:

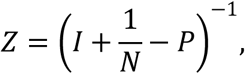

It is easy to prove that the symmetric matrix *I* + 1/*n* − P has no zero eigenvalues, ensuring the existence of *Z*. The inversion of the symmetric matrix can be efficiently performed using QR decomposition. Therefore, *M* can be solved rapidly even at high resolutions.

The MFPT matrix obtained by this algorithm has two practical limitations. First, its diagonal elements are all *n*, and most of its elements are close to *n*. Second, the matrix is asymmetric. Considering the biological significance of contact frequencies, both the ease of transition from *i* to *j* or from *j* to *i* implies the spatial proximity between bin *i* and *j*. Therefore, we adopt the following form for the MFPT representation:

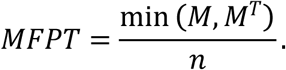

which resulting MFPT representation symmetric, with all elements strictly positive, and all diagonal elements consistently 1.

### Determination of STRIDE metric

For a pair of contact matrices with MFPT representations _1_ and _2_, their STRIDE distance is directly based on the matrix norm of the difference between the log-transformed MFPT representations, while normalizing the distance using the modulus of the average of the two matrices to ensure comparability across different chromosomes. The formula for STRIDE is as follows:

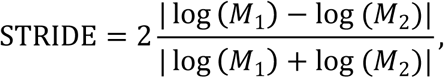

where | ∗ | denotes any matrix norm. In this paper, the 2 norm is used exclusively for all bulk Hi-C datasets and Frobenius norm is used for the single cell Hi-C datasets.

The special form of the Frobenius norm simplifies the calculation of STRIDE for a large batch of contact matrices simultaneously. Suppose there are *m* MFPT matrices (after log-transformation) to be computed. They are firstly flattened with only the upper triangular part retained, then recombined into a new matrix *X*_(*n*(*n*−1)/2)×*m*_. Based on its Gram matrix *G* = *X^T^X*, it is straightforward to compute:

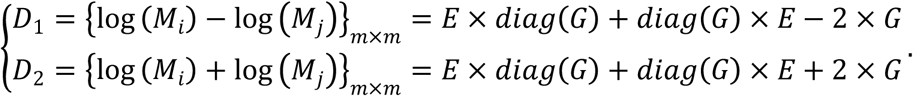

Thus, the pairwise STRIDE can be efficiently computed using the following matrix formula:

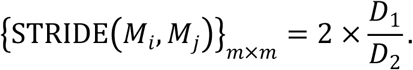

In this study, the calculation of STRIDE in bulk Hi-C libraries consistently utilizes a 50Kb resolution. This resolution effectively balances computational complexity with the ability to capture detailed differences within Hi-C maps and has been widely adopted in numerous relevant publications. For single-cell Hi-C libraries, the computation of STRIDE is performed at a 500Kb resolution, primarily due to considerations of computational feasibility.

Empirically, at a 50Kb resolution in high-quality Hi-C libraries, the STRIDE scores between biological replicates typically remain below 0.1, and even in lower-quality libraries, these scores rarely exceed 0.2. Consequently, these two thresholds can be recommended as thresholds to determine whether two libraries can be considered as biological replicates.

### Determination of TAD boundaries based on insulation scores profiles in both contact maps and MFPT maps

In both contact and MFPT maps, insulation score (IS) profiles are computed chromosome-by-chromosome following the configuration methodology in our previous work at a 10Kb resolution ^40^. A sliding window of 250Kb radius is employed for these calculations. The resulting IS profiles are normalized by their respective chromosome averages and subsequently subjected to a logarithmic transformation to facilitate comparisons of score variations at TAD boundaries and within TAD bodies across different sequencing depths.

TAD boundary annotations are derived from the contact map of the complete library, as detailed in our prior study, and are consistently applied throughout all analyses. For each chromosome, insulation score values are modeled using a three-component Gaussian mixture model, which categorizes them into strong boundaries, weak boundaries, and TAD interiors. Model parameters are estimated via the Expectation-Maximization (EM) algorithm. The classification of each bin is then determined based on the component with the highest posterior probability. Bins inferred as boundaries (whether strong or weak) that also correspond to local minima of the insulation scores within their 21-bin neighborhoods are identified as candidate boundaries. Adjacent boundaries are consolidated into TADs if more than 50% of the intervening bins are classified as weak boundaries or interiors.

### Generation of loop position annotations and application of APA analysis

The reference loop positions are identified in the complete library’s contact map at 10Kb resolution using the mustache package with default parameters ^41^, and this reference is consistently applied across all comparisons. The calculation of loop signal enrichment scores and the application of APA ^42^ analysis follow identical strategies in both the contact and MFPT maps. Let *M* represent the map under consideration. For each loop involving bins *a* and b (*a* < *b*), the average and standard deviation of signal values in submatrices *M*[(*a* − 6): (*a* − 3), (*b* − 6): (*b* − 3)] and *M*[(*a* + 3): (*a* + 6), (*b* + 3): (*b* + 6)] are computed (denoted as *m* and *s*, respectively). The loop signal enrichment score (*z* score) is then derived as:

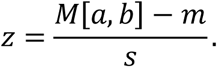

APA analysis is performed by averaging all normalized local signal maps (*M*[(*a* − 10): (*a* + 10), (*b* − 10): (*b* + 10)] − *m*)/*s* across all loops. A global *z* score is calculated as the mean of all individual loop *z* scores. To mitigate the impact of rapidly decaying contact frequencies near the diagonals of the contact maps, only loops with anchor distances ≥ 150Kb (10 bins) are included in these calculations.

In the context of these maps, a higher *z* score indicates greater significance of the loop in terms of contact frequencies for contact maps, while the interpretation is reversed for MFPT maps.

### Pseudo-replication generation and down-sampling

The pseudo-replicates of the GM12878 libraries are achieved by evenly partitioning the library into two sub-libraries. Let the raw contact map be denoted as *M*. One pseudo-replicated contact map (*M*_1_) is generated by populating its elements with random numbers drawn from a Bernoulli distribution, where *M*_1_[*i*, *j*] ∼ *B*(*M*[*i*, *j*], 0.5). The second pseudo-replicate (*M*2) is constructed as *M*_2_ = M − M_1_. Down-sampled libraries are created by dividing each contact count into batches corresponding bin pairs, merging these batches, randomly sampling a subset of bin pairs without replacement according to the desired sampling rate, and subsequently reconstructing the contact map using the selected bin pairs.

### Application of commonly used library consistency metrices

In this study, we compared the performance of three widely used Hi-C library consistency metrics: HiCRep ^18^, HiC-spector ^20^, and genomeDISCO ^21^. Although some publications highlight the performance of another metric, QuaSAR-seq ^19,43^, we exclude it from our comparison because it is tightly integrated into the hifive package ^44^ and cannot be run independently with plain text or.hic format contact maps as inputs.

All metrics, including STRIDE, are analyzed chromosome by chromosome. We do not average scores across all chromosomes to construct a uniform score like most previous publications do, as scores vary significantly among chromosomes.

The HiCRep package is sourced from its GitHub repository (https://github.com/TaoYang-dev/hicrep) rather than Bioconductor. In our application, the maximum genome distance allowed in the similarity score calculation is set to infinity to ensure the algorithm considers the full map, aligning it with the other three metrics. The parameter ℎ, which controls the smoothing level, is fixed at 5 for all sequencing levels and chromosomes. This choice is due to the computational intensity required to determine this value and the recommendation by the authors for a resolution of 40Kb, which is close to the resolution used in this study. The feasibility of this choice has been validated by determining optimal values in several chromosomes.

The source code of HiC-spector is obtained from the GitHub repository associated with the original publication (https://github.com/gersteinlab/HiC-spector) and executed using default configurations. The number of eigenvectors taken is set to 20, as recommended.

A private implementation of genomeDISCO, compatible with Python3 and parallel processing, is utilized. The number of transitions is set to 3, following the publication’s recommendation, and the normalization type is specified as “sqrtvc.” Both HiCRep and genomeDISCO offer options to balance sequencing depths by down-sampling deeper-sequenced libraries. These options are consistently enabled. In contrast, HiC-spector lacks this feature. We do not artificially introduce such functionality and allow the algorithm to operate on the original contact maps.

The final steps of the three metrics—HiCRep, HiC-spector, and genomeDISCO—involve calculating a similarity score analogous to the Pearson correlation coefficient. These scores are converted into pairwise distances using straightforward linear algebra. If the similarity score matrix is denoted as *C*, the corresponding distance matrix *D* is derived as 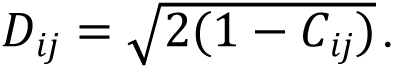 All hierarchical clustering and calculations of CH indices are performed on the distance matrix rather than the original similarity matrix.

## Data availability

All datasets used in this study are publicly available. Hi-C data of individual and merged GM12878 cells and the merged K562 cells are downloaded from the Gene Expression Omnibus (GEO) database under accession number GSE63525. The Hi-C replicates of hESC and IMR90 cells derived using HindIII are downloaded under accession number GSE35156. The replicates from hESC derived from DpnII are downloaded under accession number GSE165894. The replicates of IMR90 derived from MboI are downloaded under accession number GSE63525. Hi-C and RNA-seq data in mouse hematopoietic system are downloaded under accession number GSE152918 and GSE142216, respectively. Hi-C data in early porcine embryo development are downloaded under accession number GSE153452. Single cell Hi-C data in the mESC cell cycle are download under accession number GSE94489. The mouse cortical and hippocampal neurons are downloaded under the accession number GSE162511. The details of libraries involved are listed in (Extended Data Table 1).

## Acknowledgements

This work was supported by the National Natural Science Foundation of China (32200515 to XBX, 32470672 to LFF and 32341011 to ZZH), the Strategic Priority Research Program of CAS (XDA0460300, XDC0200000 to ZZH), Biological Breeding-National Science and Technology Major Project (2022ZD04017 to ZZH). We would like to thank Professor Ru Wang from Shanghai University of Sport for help in the computational resources and Professor Feng Liu from Hebei University of Technology for their valuable suggestions.

## Author Contributions

XBX and ZZH conceived this project. XBX and LFF analyzed data. XBX, LFF, TWC and ZZH prepared the manuscript. GXM prepared the figures and tables.

## Competing interests

The authors declare no competing interests.

## Code availability

The code and data which generated the results of this study are stored in gitee and freely accessible at the URL https://gitee.com/matrix_evolution/STRIDE. The STRIDE package can also be freely available at PyPi following the URL https://pypi.org/project/hicstride.

## Notes

### Competing Interest Statement

The authors have declared no competing interest.

