## supplemental text and figures for "STRIDE: A Sequencing Depth-Insensitive Metric for Robust Comparison between Sparse Chromosome Conformation Capture Data"

### Public data analysis.

All public Hi-C datasets employed in this study are enumerated in (Extended Data Table 1). Each downloaded read underwent rigorous quality assessment via FastQC (accessible at https://github.com/s-andrews/FastQC). Reads exhibiting substandard sequencing quality (quality score $\leq10$) and those containing adaptor sequences were meticulously purged utilizing the fastp tool. Subsequently, the refined reads were channeled through the HiC-Pro pipeline, adopting its default parameters for processing. To maintain consistency with their source publications, the human reference genome hg19 was uniformly applied to the GM12878, K562, hESC, and IMR90 datasets.

For matrix equilibration, the Kilian-Rajagopalan (KR) algorithm was employed. Initially, all diagonal entries within the contact matrix were fitted by zeros. This step preceded the exclusion of bins characterized by an excessively low depth of sequencing coverage, specifically targeting those whose row aggregates did not surpass the second percentile of row sums from bins with positive row sums. Upon reaching convergence, an additional refinement was instituted to ensure that both row and column summations were rigorously normalized to a unit value. Let $s_{i}$ represent the row sum of bin $i$ in the balanced matrix $\{w_{ij}\}$, and let $m$ denote the maximum among all $s_{i}$ values. The elements $w_{ij}$ were then adjusted according to the formula:

$$w_{ij}=\left\{ \begin{aligned} w_{ij}/m, &i\neq j \\ 1-s_{i}/m, &i=j \end{aligned} \right..$$

### Calculating the signal to noise of contact frequencies or MFPT maps.

The signal-to-noise ratio (SNR) was computed based on either KR-normalized contact frequencies or mean first passage time (MFPT) values. In this context, data from fully sequenced libraries were considered as the 'clean' signals, while those from down-sampled libraries represented signals contaminated with noise. The analysis was conducted on maps with a resolution of 10 kilobases (Kb). Only genomic regions with distances not exceeding 2 megabases (Mb) were included in the analysis. For contact frequency calculations, regions exhibiting zero values in either the fully sequenced or down-sampled libraries were preemptively excluded. Subsequently, the values were subjected to a log2 transformation. A linear regression analysis was then performed, wherein the transformed values from the down-sampled libraries were regressed against those from the fully sequenced libraries. The SNR was subsequently defined as the ratio of the regression sum of squares to the residual sum of squares derived from the regression model. This computation was executed separately for each chromosome.

### Calculating the genome distance for the variability of contact frequencies (MFPTs) to stabilize.

For a (chromosome-wise) contact map or MFPT map at 10Kb resolution $\left\{ c_{ij} \right\}_{n\times n}$, the coefficient of variance (CV) of the $\log_{2}$ transformed contact frequencies or MFPT values along the same genome distances were determined as

$$cv_{d}=\frac{std(\{c_{ij}|j-i=d\})}{mean(\{c_{ij}|j-i=d\})},d=1,2,\cdots300.$$

Then, the genomic distance at which the CV values reached their stability was determined as the abscissa of the point where the curve $cv(d)$ tangent to the line with slope $k=\frac{cv\left( 300 \right)-cv(1)}{299}$ among all lines with that slope.

### Calculation of the intra- and inter- TAD average contact frequencies or MFPTs around TAD boundaries.

For a contact frequency or MFPT map at 10Kb resolution $\left\{ c_{ij} \right\}_{n\times n}$ and a TAD boundary $i$, the intra- and inter-TAD average contact frequency or MFPT was calculated as follows:

$$\left\{ \begin{aligned} \mathrm{intra}=\frac{1}{1640}(\sum_{k=i-50}^{i-5} \sum_{l=k+5}^{i-5} c_{kl}+\sum_{k=i+5}^{i+45} \sum_{l=k+5}^{i+50} c_{kl}) \\ \mathrm{inter}=\frac{1}{3016}(\sum_{k=i-50}^{i+45} \sum_{l=k+5}^{i+50} c_{kl}-\mathrm{intra}) \end{aligned} \right..$$

### Sequencing depth sensitivity index and discrimination index calculation for a library (in)consistency metric.

Suppose in the series of downsampling procedure, the function of consistency scores against the $\log_{2}$ transformed sampling rate for PR, BR and NR were denoted as $p\left( d \right), b\left( d \right)$ and $n\left( d \right)$ where $d=0,1,2,\cdots,5$. Scores were firstly transformed into the $\left[ 0,1 \right]\times\left[ 0,1 \right]$ square by taking the following transform

$$f^{'}\left( d^{'} \right)=f(5\times d)/max(n\left( 5 \right),$$

where $f$ represented one of $p$, $b$ or $n$, $d^{'}$=$\frac{d}{5}$. Then, each of these functions was fitted with a 5th-order polynomial where all coefficients were non-negative. The results of the fitting were still denoted as $p^{'}$, $b^{'}$ or $n^{'}$. The sequencing depth sensitivity index was then calculated as

$$\int_{0}^{1} f^{'}\left( t \right)dt-f\left( 0 \right),$$

where $f=p, b \mathrm{or} n$.

The discrimination indexes between PR and BR, as well as between BR and NR, were calculated as the roots of the equations $p^{'}\left( t \right)=b\left( 0 \right)$ and $b^{'}\left( t \right)=n\left( 0 \right)$, respectively.

### Extended Data Figures

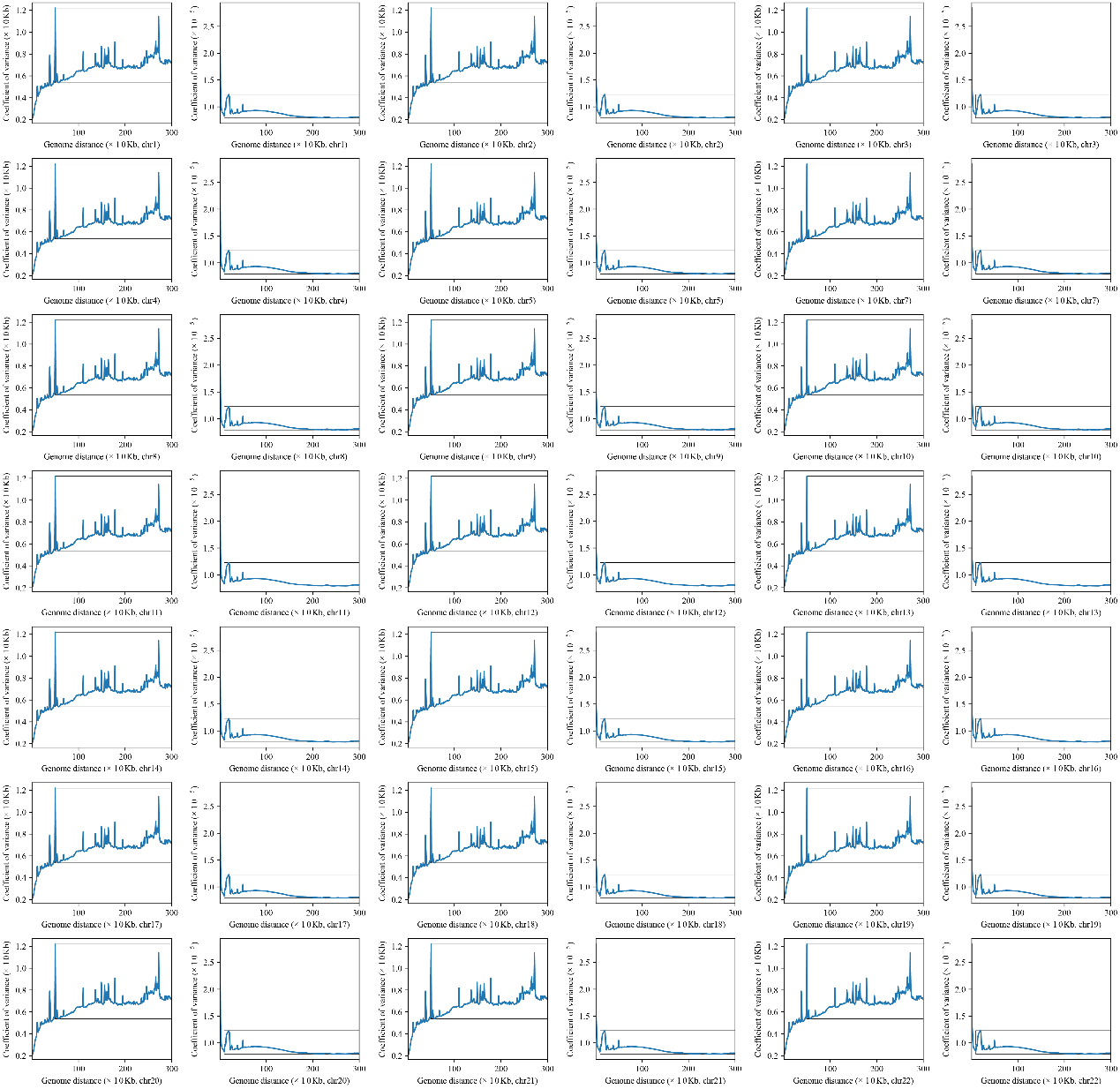

**Extended Data Fig. 1 The relationship between the variability (coefficient of variation, CV) of contact frequencies (left panels) or MFPT values (right panels) and genomic distances in chromosomes other than chromosome 6 (which have been shown in Fig. 1E and F).** For the contact maps, zeros were removed from the calculation of the CV. The markers were the same as those in Fig. 1E and F.

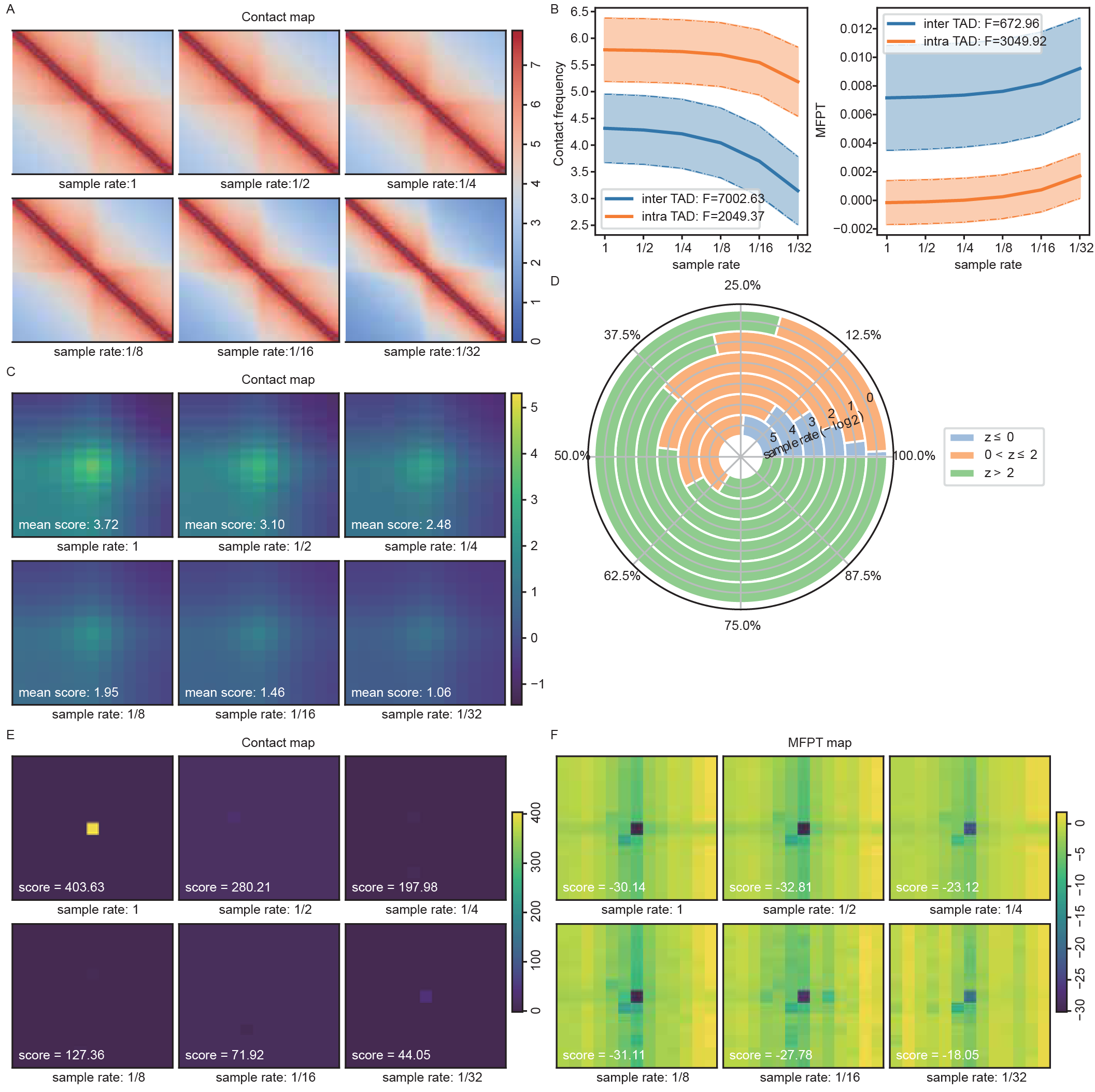

**Extended Data Fig. 2 MFPT representation preserves topological features in low-depth Hi-C libraries than contact map.** **(A)** The average contact profiles around annotated TAD boundaries in contact maps in all sample rates. **(B)** The mean (solid lines) and standard deviations (shades) of the average intra- and inter-TAD contact frequencies or MFPT distances around TAD boundaries in libraries with different sequencing depths. The AUC scores, derived from the areas between curves and dashed lines, measuring the sensitivity of the frequencies or distances to the sequencing depths were also shown (Supplementary text). The larger the value, the more sensitive of the frequencies or distances to the decay of sequencing depth. **(C)** The APA maps calculated from the contact maps in all sample rates, with average $z$ scores marked. **(D)** Proportions of annotated loops with significantly enriched contact signals ($z>2$), with enriched but not significant contact signals ($0\leq z<2$), and with no enriched contact signals ($z\leq0$). The $z$ scores were calculated from contact maps. **(E)** and **(F)** The normalized contact profiles around the loop between chr22:22.97Mb and 24.65Mb calculated from contact maps (D) and MFPT maps (E) in all sample rate, with the $z$ scores of the loop marked.

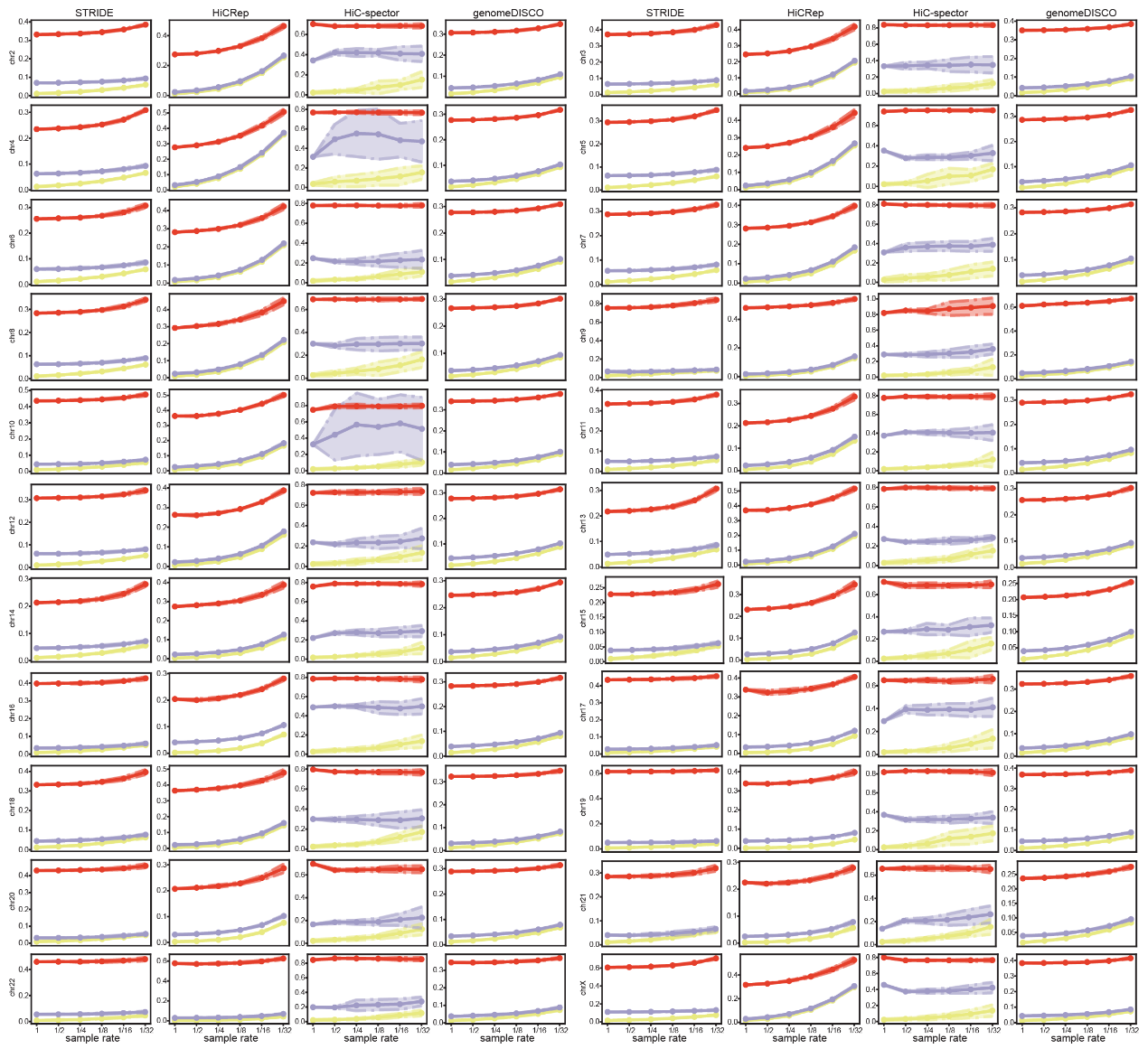

**Extended Data Fig. 3 The similarity and dissimilarity scores computed by the four metrics among PRs, BRs, and NRs at each sampling rate in chromosomes other than chr1.**

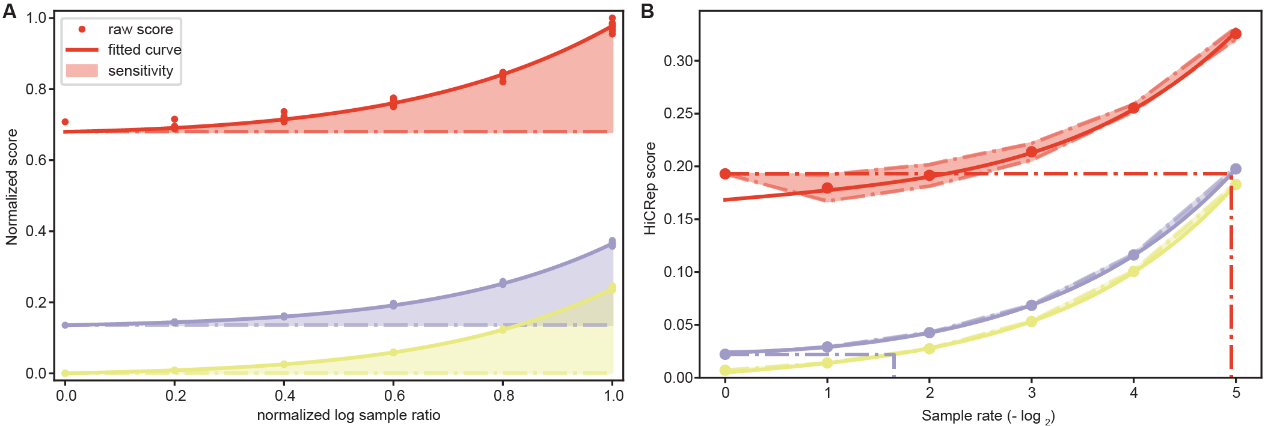

**Extended Data Fig. 4 Schematic diagram for calculating sequencing depth sensitivity and discrimination index among PR, BR, and NR. (A)** HiCRep of chr1 was shown as schematic diagram for calculating sequencing depth sensitivity. **(B)** HiCRep on chr16 was shown as schematic diagram for discrimination index among PR, BR, and NR.

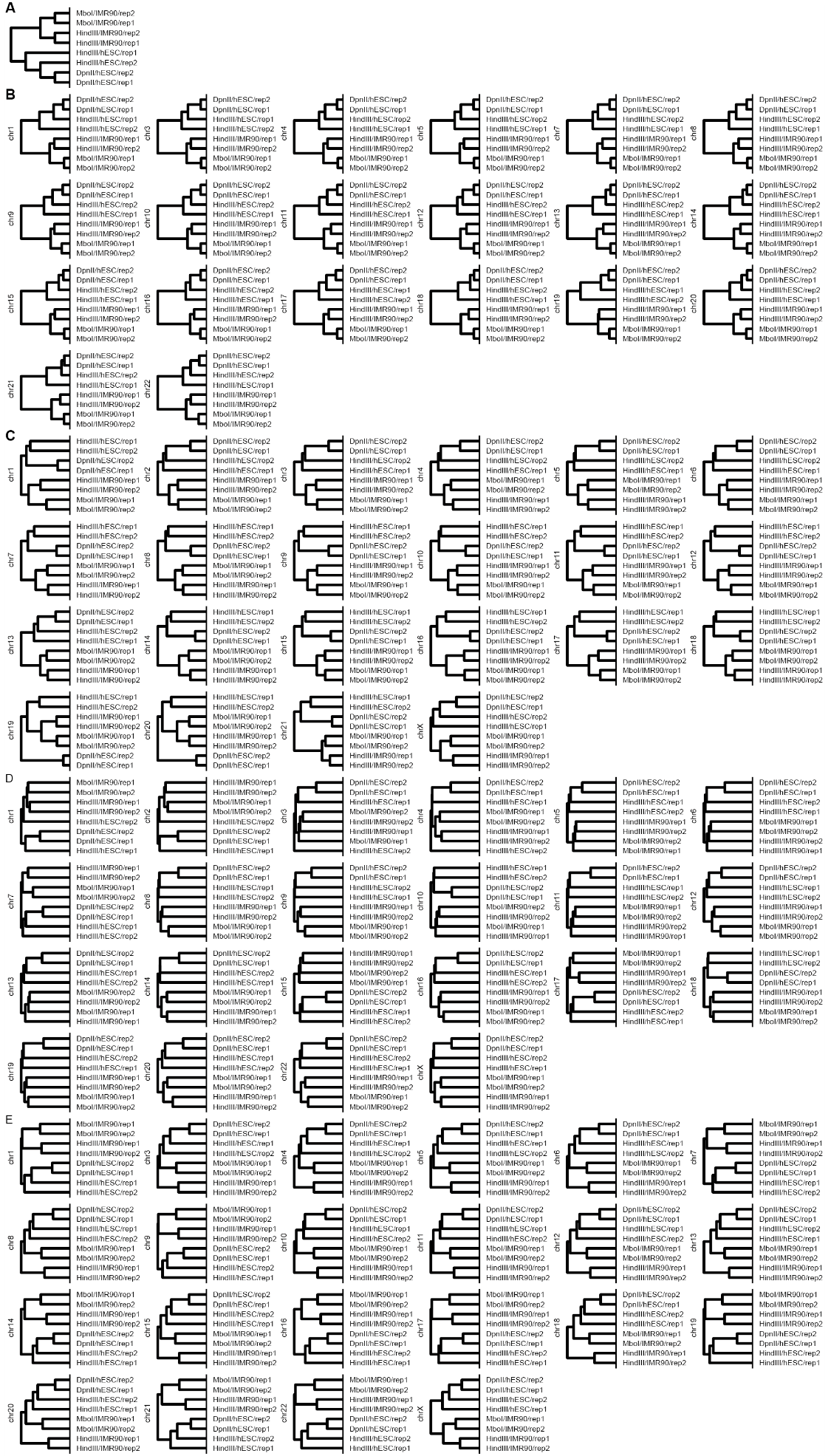

**Extended Data Fig. 5 Hierarchical clustering results of the IMR90 and hESC libraries calculated from library consistencies or distances in chromosomes not included in Fig. 4B.** The expected clustering hierarchy of the 8 libraries was shown in (A). Resulted clustering hierarchies from STRIDE, HiCRep, HiC-spector and genomeDISCO were shown in (B), (C), (D) and (E), respectively.

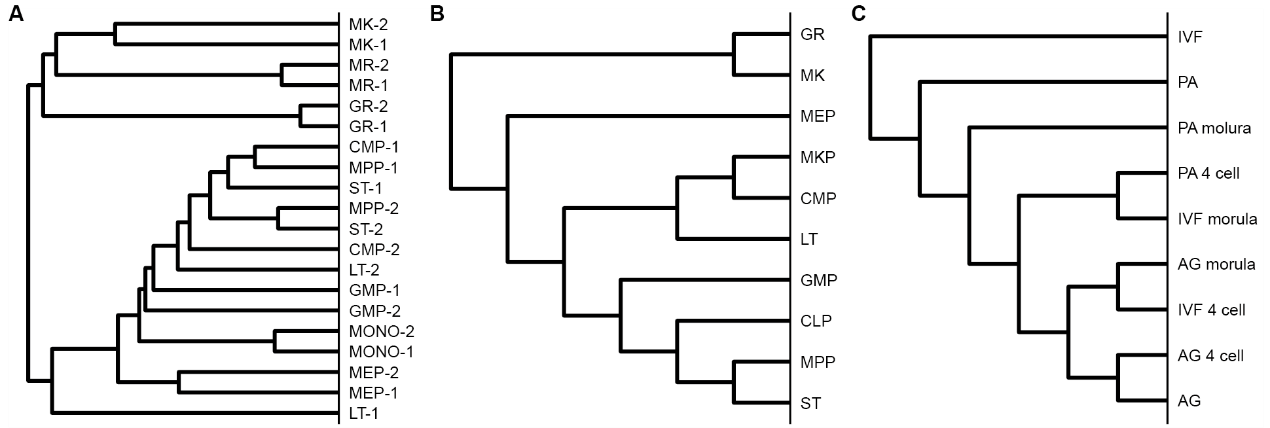

**Extended Data Fig. 6 The reference phylogenetic relationships among cell types. (A)** The hierarchical clusters of the cells in mouse hematopoietic system generate by RNA-seq. **(B)** The reference phylogenetic relationship among cells in mouse hematopoietic system. **(C)** The reference phylogenetic relationship among cells in early porcine embryo development.

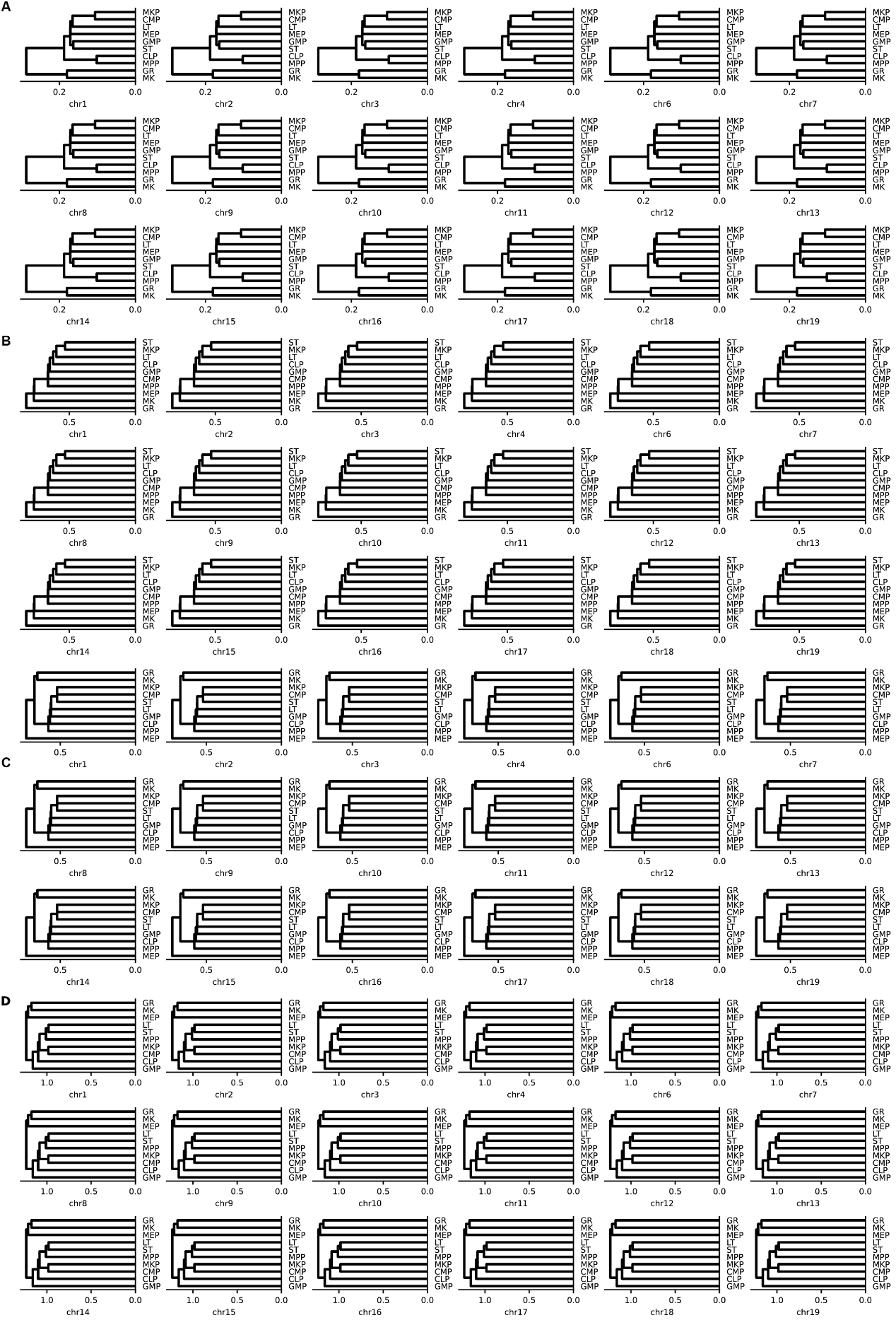

**Extended Data Fig. 7** **Hierarchical clustering dendrograms of cell types in mouse hematopoietic system in chromosomes other than chr5.** Results from STRIDE, HiCRep, genomeDISCO and HiC-spector were shown in (A), (B), (C) and (D), respectively.

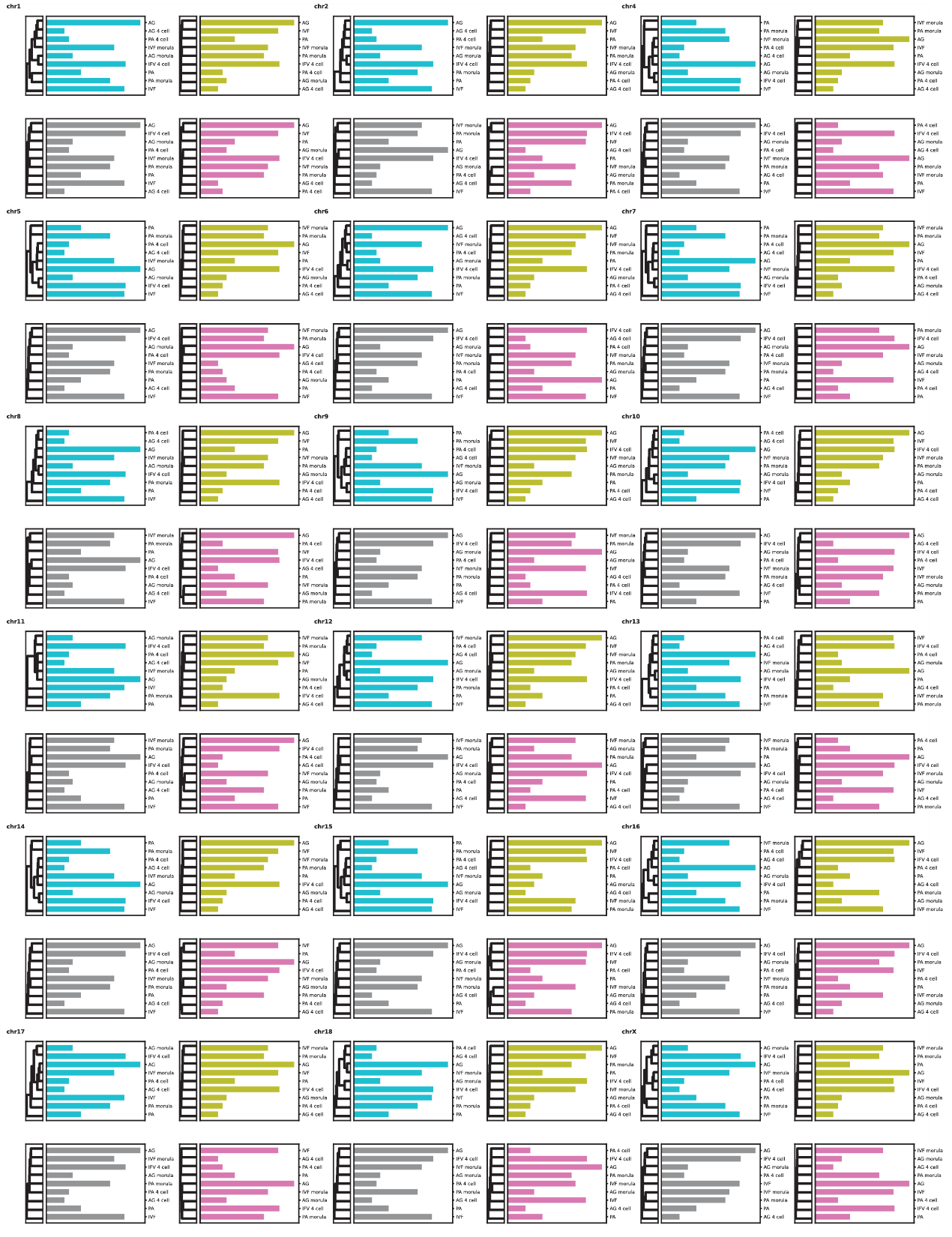

**Extended Data Fig. 8 The hierarchical clustering results among cells in early porcine embryo development in chromosomes other than chr3.** The widths of the bars marked the sequencing depths. The color code was the same as in Fig. 5C.

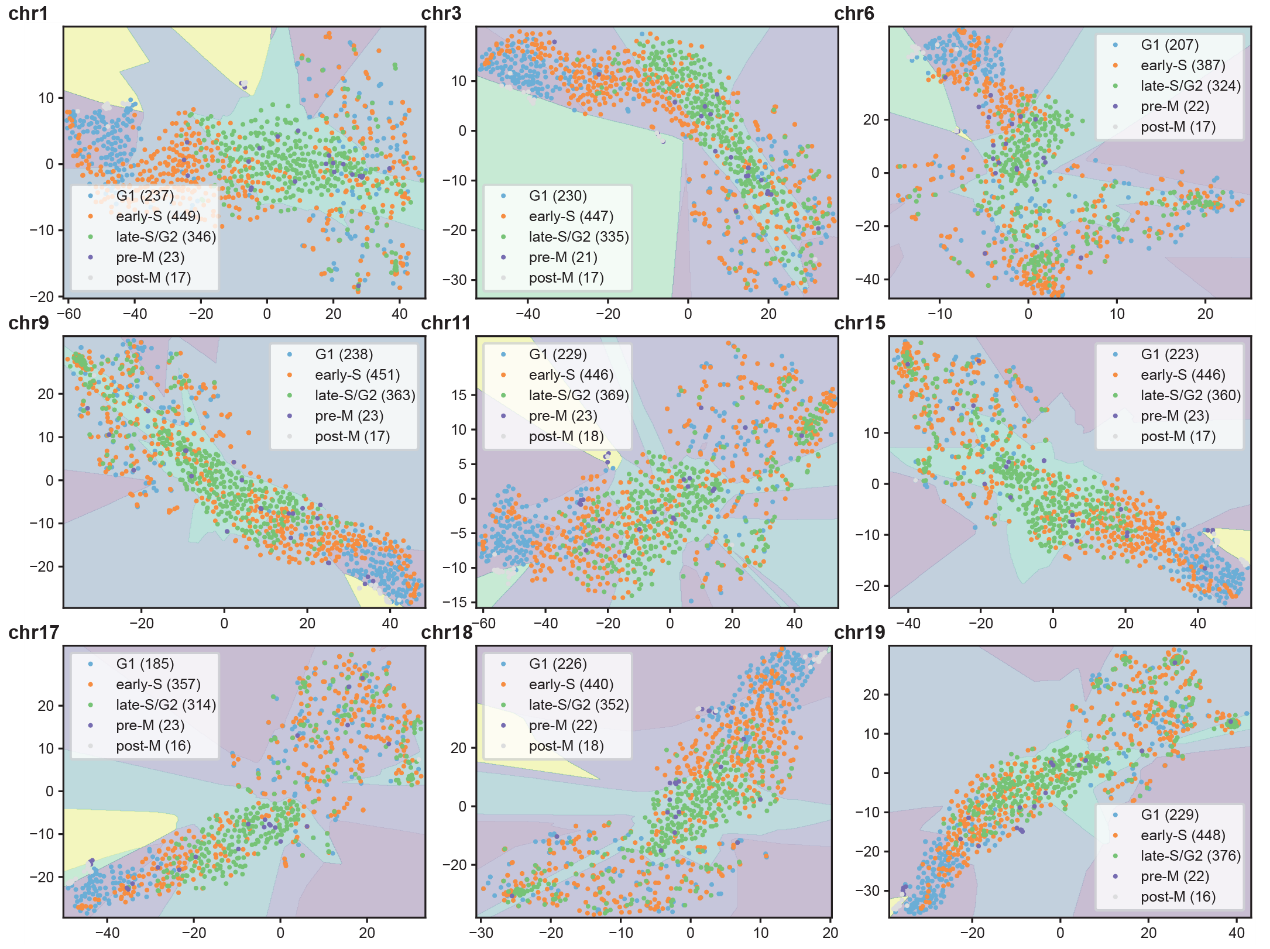

**Extended Data Fig. 9 The tSNE low-dimensional embedding results on the cell cycle dataset in chromosomes other than that shown in Fig. 6A.** The shaded areas represent the cell type separation results based on the MLP model.

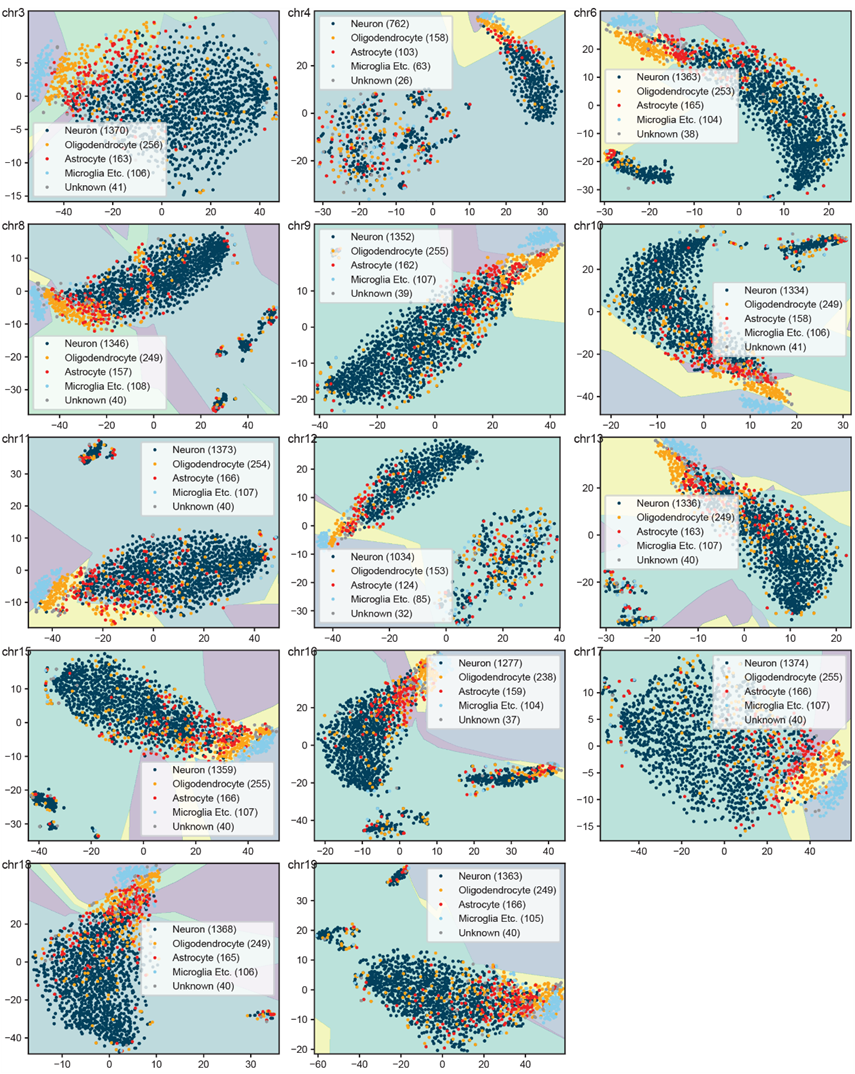

**Extended Data Fig. 10 The tSNE low-dimensional embedding results on the mouse neuron dataset in chromosomes other than that shown in Fig. 6B.** The shaded areas represent the cell type separation results based on the MLP model.

| **Dataset name** | **Accession number** | **Source** | **Files or libraries used** |
| --- | --- | --- | --- |
| **Hi-C libraries** |  |  |  |
| Hi-C on GM12878 and K562 | GSE63525 | 10.1016/j.cell.2014.11.021 | All libraries from GM12878 or K562, merged into primary or replicate as the Supplementary Table 1 in the publication noted. |
| Original Hi-C libraries of hESC and IMR90 | GSE35156 | 10.1038/nature11082 | GSM862720, GSM862721, GSM862724, GSM892307 |
| The Hi-C libraries from IMR90 using MboI | GSE63525 | 10.1016/j.cell.2014.11.021 | GSM1551604, GSM1551605 |
| The Hi-C libraries from hESC using DpnII | GSE163666 | 10.1038/s41592-021-01248-7 | GSM5057434, GSM5057435 |
| Hi-C libraries in mouse hematopoiesis | GSE152918 | 10.1016/j.celrep.2020.108206 | All Hi-C libraries |
| Hi-C libraries from early porcine embryo | GSE153452 | 10.1186/s13059-020-02095-z | All Hi-C libraries |
| **polyA+ RNA-seq datasets** |  |  |  |
| RNA-seq libraries in mouse hematopoiesis | GSE142216 | 10.1016/j.celrep.2020.108206 | Libraries from the 8 cell types which conicide with the Hi-C libraries. |
| **Single cell Hi-C libraries** |  |  |  |
| Single cell Hi-C libraries in mESC cell cycle | GSE94489 | 10.1038/nature23001 | All samples passing the quality thresholds mentioned in the publication together with the sample meta data. |
| Single cell Hi-C libraries mouse cortical and hippocampal neurons | GSE162511 | 10.1016/j.cell.2020.12.032. Epub 2021 Jan 22 | All samples passing the quality thresholds mentioned in the publication together with the sample meta data. |

**Table S1. Public dataset used in this study**
